# Antagonistic effects of arginine methylation of LSM4 on alternative splicing during plant stress responses

**DOI:** 10.1101/2023.12.08.570794

**Authors:** Yamila Carla Agrofoglio, María José Iglesias, Soledad Perez-Santángelo, María José de Leone, Tino Koester, Rafael Catalá, Julio Salinas, Marcelo J. Yanovsky, Dorothee Staiger, Julieta L. Mateos

**Affiliations:** Instituto de Fisiología, Biología Molecular y Neurociencias (IFIBYNE-UBA-CONICET) and Facultad de Ciencias Exactas y Naturales, Universidad de Buenos Aires, Ciudad Universitaria, Buenos Aires, Argentina; Fundación Instituto Leloir, Instituto de Investigaciones Bioquímicas de Buenos Aires–Consejo Nacional de Investigaciones Científicas y Técnicas (CONICET), C1405BWE Buenos Aires, Argentina; RNA Biology and Molecular Physiology, Faculty of Biology, Bielefeld University, 33615 Bielefeld, Germany; Departamento de Biotecnología Microbiana y de Plantas, Centro de Investigaciones Biológicas Margarita Salas, CSIC, 28040 Madrid, Spain

**Author notes:** Department of Biochemistry, University of Otago, Dunedin, 9016, New Zealand (Soledad Perez-Santángelo).

**Keywords:** list *6–10 keywords*, arginine methylation, alternative splicing, LSM4, PRMT5, Arabidopsis, abiotic stress, absicic acid, biotic stress

## Abstract

Arabidopsis PROTEIN ARGININE METHYLTRANSFERASE 5 (PRMT5) post-translationally modifies RNA-binding proteins by arginine (R) methylation. The impact of this modification on the regulation of RNA processing is largely unknown. Here we use LSM4, a component of the spliceosome, as a paradigm to study the impact of R-methylation on its function in RNA processing. We identify in vivo targets of LSM4 and show that LSM4 regulates alternative splicing of a suite of them. Furthermore, LSM4 affects mRNA levels of some of the targets, showing for the first time its role in both AS and steady-state abundance. The *lsm4* and *prmt5* mutants show a considerable overlap of genes with altered splicing patterns, suggesting that these might be regulated by PRMT5-dependent LSM4 methylation. Wild-type LSM4 and an unmethylable version complement the *lsm4-1* growth and circadian rhythms defects, suggesting that methylation is not critical for growth in normal environments. However, LSM4 methylation increases with ABA and is necessary for plants to respond properly to salt stress. In contrast, LSM4 methylation is reduced by bacterial infection, and plants expressing unmethylable LSM4 are more resistant than plants expressing wild-type LSM4. This tolerance correlates with decreased intron retention of immune-response genes upon infection, augmenting the functional isoform. Taken together, this provides the first direct evidence that R methylation adjusts LSM4 function on pre-mRNA splicing in an antagonistic manner in response to biotic and abiotic stress.

**Highlight:** *Please provide a statement that, in fewer than 30 words, highlights the novelty of the paper for the non-expert*.

Arginine methylation of the LSM4 spliceosome component by PROTEIN ARGININE METHYLTRANSFERASE 5 fine-tunes alternative splicing of a set of stress-related genes to antagonistically control biotic and abiotic responses in Arabidopsis.

## Introduction

The ability of plants to acclimatise and maintain yield under stressful environmental conditions is a topic of heightened concern due to climate change and increasing demands on agriculture. Plant stress acclimation involves multiple layers of post-transcriptional (Hernando *et al*., 2017; Dikaya *et al*., 2021) and post-translational regulation (Qin *et al*., 2008; Withers and Dong, 2017; Benlloch and Lois, 2018; Augustine and Vierstra, 2018). Pre-mRNA splicing is crucial for adequate plant stress responses, and while splicing is greatly influenced by different stresses (Staiger and Brown, 2013; Laloum *et al*., 2018; Rigo *et al*., 2019), little is known about how stress signalling affects the splicing machinery to mediate such changes in splicing.

During pre-mRNA splicing, removal of introns is accomplished by the spliceosome, a high molecular weight complex consisting of five small nuclear ribonucleoprotein particles (snRNPs) and over 200 additional proteins (Wahl *et al*., 2009). The snRNPs aid in recognition of the 5’ and 3’splice sites and contribute to the catalytic mechanism of the spliceosome. The five snRNPs contain the name-sake small nuclear uridine-rich RNAs (U1, U2, U4, U5, and U6 snRNAs). The core proteins of the U1, U2, U4, and U5 snRNPs are the Sm proteins, firstly found with antibodies from a serum lupus erythematosus patient (Stephanie Smith (Sm)). In contrast, the U6 snRNP contains the related LSM (Like Sm) proteins 2 to 8. All eukaryotes have seven highly conserved Sm proteins (B/B’, D1, D2, D3, E, F, and G) and eight LSM proteins that typically exist as hexameric or heptameric complexes in vivo. In mammalian cells, Arginine (R) methylation of Sm B/B′, D1, and D3 proteins is essential for snRNP assembly (Wahl *et al*., 2009). Still, in Drosophila, methylation of Sm proteins is not critical for snRNP biogenesis, highlighting species-specific differences in this basal cellular process (Gonsalvez *et al*., 2008)

Sm and LSM R methylation is catalysed by Protein Arginine Methyltransferases (PRMTs) which transfer the methyl group from S-adenosylmethionine to the guanidinium nitrogen atoms of the R residue. PRMTs are conserved from plants to humans and the importance of R methylation is underscored by impaired PRMT activity associated with autoimmune diseases or cancer in mammals. PRMTs are divided into three types, based on the kind of methylation they promote. Type I, which comprises PRMT1-4, 6 and 8, promotes monomethylation and asymmetric dimethylation of arginines (MMA and ADMA, respectively). Type II, which includes PRMT5 and 9, promotes MMA and symmetric dimethylation of arginines (sDMA). And last, type III, which consists of only PRMT7 promotes MMA of arginines (Bedford, 2007).

In Arabidopsis, the best characterised PRMT is the type II methyltransferase PRMT5. Mutants of *PRMT5* are late flowering, have altered circadian rhythms (Wang *et al*., 2007; Pei *et al*., 2007; Sanchez *et al*., 2010; Hong *et al*., 2010), are more susceptible to various abiotic stresses (Zhang *et al*., 2011; Hernando *et al*., 2015; Hu *et al*., 2017) and have widespread defects in the splicing of thousands of genes (Deng *et al*., 2010; Sanchez *et al*., 2010; Hernando *et al*., 2015). Whether this is due to its role as a methyltransferase remains to be elucidated.

Over the years, PRMT5 targets have been identified in plants, including the U snRNP proteins SmD1, SmD3 and LSM4, and the RNA-binding protein *At*GRP7, all involved in splicing (Deng *et al*., 2010; Hu *et al*., 2019; Cao *et al*., 2022, Streitner *et al*., 2012; Wang *et al*., 2020) suggesting that the R methylation of splicing factors could provide an efficient means to fine-tune their activity. This raised the idea that PRMT5 may affect alternative splicing (AS) of some pre-mRNAs indirectly by modulating the activity of spliceosome components through R methylation. But so far, not much is known about the direct role of R methylation in AS. The *lsm4* mutant plants show severe growth retardation, are sterile and their response to salt stress and abscisic acid (ABA) is found to be severely impaired (Zhang *et al*., 2011). Moreover, LSM4 affects circadian clock function, likely by regulating the AS of an unknown set of core clock genes, contributing to adjustment to periodic changes in environmental conditions (Perez-Santangelo *et al*., 2014). Recently, LSM4 has also been implicated in the regulation of plant immunity to *Pseudomonas syringae* infection by interacting with the metacaspase AtMC1, modulating AS of genes such as *4-COUMARATE:COA LIGASE 3* (*4CL3*), a known negative regulator of plant defense (Wang *et al*., 2021).

In this study, we used LSM4 as a paradigm to investigate the effect of methylation of R by PRMT5 on AS. Through RNA immunoprecipitation, we identified transcripts bound by LSM4 in vivo and we show that for a suite of these targets, LSM4 regulates AS by directly binding to its transcripts. Furthermore, mRNA levels of direct targets were also affected by LSM4 binding, showing for the first time the direct role of LSM4 in both AS and steady-state mRNA levels. In addition, the *lsm4* and *prmt5* mutants showed a significantly high overlap of genes with altered splicing patterns, suggesting that these might be regulated by PRMT5-dependent methylation of LSM4. Using complementation studies, we show that a wild-type version, as well as an unmethylable version of LSM4, were able to rescue the *lsm4-1* mutant under controlled growth conditions, suggesting that methylation is not critical for plants growing in normal growth conditions. However, LSM4 methylation changes along with stress-related environmental cues. LSM4 methylation increases with ABA and is necessary for plants to respond properly to salt stress. In contrast, LSM4 methylation is reduced by bacterial infection, and plants expressing a unmethylable LSM4 show a better immune response than plants expressing the wild-type version of LSM4. Interestingly, this phenotypes correlate with our findings that, upon infection, intron retention (IR) of stress-related genes increases, diminishing the functional isoform. These data is consistent with the opposite performance of *prmt5-5* and *lsm4-1* mutants under biotic and abiotic stress.

Our results uncover a post-translational modification, R methylation, that can regulate a key post-transcriptional process, such as mRNA splicing, thus adding an additional layer of regulation to fine-tune antagonistic plant responses to bacterial infection and salt stress.

## Results

### Identification of in vivo target transcripts of LSM4

PRMT5 controls a wide range of alternative splicing (AS) events in plants (Deng *et al*., 2010; Sanchez *et al*., 2010; Hong *et al*., 2010; Hernando *et al*., 2015) and direct targets of PRMT5 methylation are enriched in RNA-binding proteins including splicing related factors such as LSM4 and *At*GRP7 (Deng *et al*., 2010; Hu *et al*., 2019, Streitner *et al*., 2012; Wang *et al*., 2020). In particular, LSM4 is part of two heteroheptameric complexes, of which the LSM1-7 complex mainly regulates transcript stability in the cytoplasm, and the LSM2-8 nuclear spliceosome complex is involved in pre-mRNA splicing through U6 snRNP (Perea-Resa et al., 2012; Golisz *et al*., 2013). But the functional relevance of R methylation on the role of LSM4 in both processes has not yet been described. As a first step to comprehensively understand the direct effects of LSM4 on mRNAs, we determined the transcripts bound by LSM4 in vivo. RNA immunoprecipitation followed by high-throughput sequencing (RIP-seq) was performed on plants expressing LSM4 tagged with YELLOW FLUORESCENT PROTEIN (YFP) under the control of the constitutive 35S promoter (35S: YFP-LSM4) in the *lsm4-1* loss-of-function mutant.

To ensure that the YFP tag does not affect LSM4 function, we confirmed the complementation of the *lsm4-1* developmental phenotype (Figure 1A). Twelve-day-old plants were crosslinked with formaldehyde to preserve in vivo RNA-protein interactions. The YFP-LSM4 fusion protein was precipitated with GFP Trap® beads and co-precipitated RNAs were sequenced. We found 982 genes to be enriched in 35S:YFP-LSM4 expressing plants relative to control plants expressing 35S:YFP only (log2 fold change > |2| and FDR < 0.05), considered from now on as LSM4 targets (Figure 1B, Table S1). Of these, we found 23 genes encoding for proteins directly involved in splicing, such as U1-70K, SMD3, SKIP2 and DRT111/SFPS, and many other RNA-binding proteins like ALBA5 and ALBA6 (Figure 1B). Consistently, we found an enrichment in Go terms, such as RNA processing, RNA splicing, and spliceosomal snRNP asswmbly for the direct LSM4 targets (Figure 1C). Furthermore, in line with the reported phenotypes for *lsm4-1* mutants (Zhang et al., 2011; Perez-Santangelo et al., 2014) LSM4 targets are enriched in terms related with ABA response, circadian rhythms and light responses (Figure 1C, Table S1).

**FIGURE 1.**
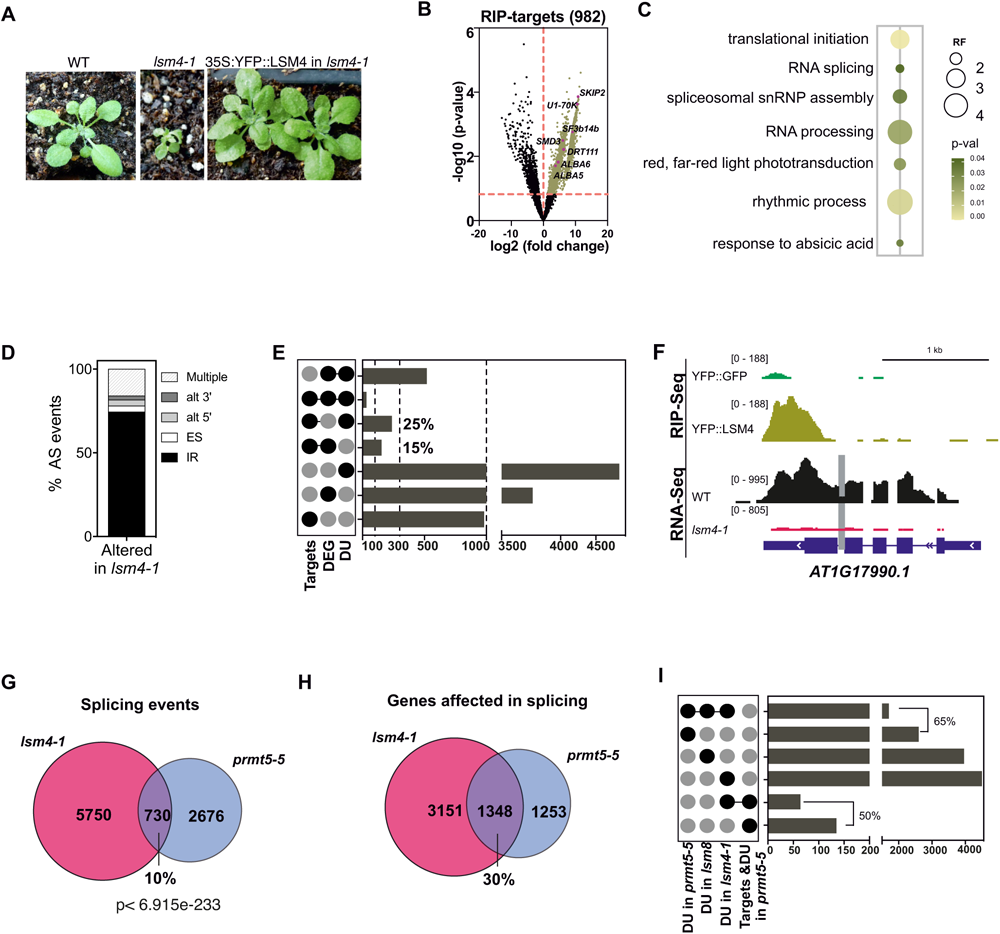
LSM4 and PRMT5 control an overlapping set of pre-mRNA splicing events. (A) Representative photograph of phenotype from plants used for RIP-Seq expressing 35S:YFP:LSM4 and *lsm4-1* and wild-type genotypes illustrating rescue of developmental defects observed in the *lsm4-1* mutant. (B) Volcano plot illustrating log_2_ fold-change (x-axis) and statistical significance as -log_10_ p-value (y-axis) of the RIP-seq dataset. The dashed line indicates the threshold above which transcripts are significantly enriched (FC >3 and p-value <0.05). (C) GO-term enrichment analysis of LSM4 direct targets represented as a bubble plot. The size of the bubble is the representation factor (RF). (D) Relative frequencies of different alternative splicing (AS) types for all detected AS events affected in *lsm4-1*: Alt 3’ and Alt 5’, alternative acceptor and donor splice sites, respectively; ES, exon skipping; IR, intron retention; Multiple, for cases where more than one type of event was observed. (E) Upset plot showing single, pairwise and triple combinations of genes for the different list of genes found in each high-throughput experiment (DEG: differentially expressed; DU: Differential usage of splicing sites from RNA-seq experiment; and Targets: Candidate genes bound to LSM4 found by RIP-Seq). (F) IGV view of mapped reads for selected LSM4-bound target transcripts in 35S:GFP and 35S:YFP:LSM4 for the RIP-seq experiment and its expression in WT and *lsm4-1* from RNA-seq. The grey line denotes the splicing defect as quantified by RNA-seq analysis. (G-H) Venn diagram showing the extent of overlap for pre-mRNA splicing events (G) or genes with splicing events (H) affected in the *lsm4-1* mutant and those altered in *prmt5-5*. (I) Upset plot of several single, pairwise and triple comparisons of gene lists from diverse analyses.

### Impact of LSM4 binding on mRNA metabolism

To address the impact of LSM4 binding on its targets RNAs, we next compared if LSM4 targets transcripts were either differentially expressed or differentially spliced in *lsm4-1* mutants. We performed RNA-seq in three biological replicates from *lsm4-1* seedlings grown for 12 days under continuous light to avoid differences caused by changes in the phase of expression due to the circadian clock defect of the mutant (Perez-Santangelo *et al*., 2014). After mapping the reads to the Arabidopsis genome, we evaluated the splicing patterns by determining the percentage of inclusion of exons (percentage splice-in, PSI) or introns (percentage intron retention, PIR) as previously described in Mancini *et al*., (2021). When comparing splicing events that differ between *lsm4-1* and wild-type plants (DU, Differential Usage), we obtained 6480 altered splicing events. These correspond to 4499 genes (Table S2), meaning that for a subset of genes, multiple splicing events were affected. We then calculate the relative distribution of the different types of AS events: alternative usage of 5’and 3’splice sites (alt 5’; alt 3’), intron retention (IR); exon skipping (ES) and multiple when more than one type was present. As reported for other splicing related proteins, IR events, the most common AS event in Arabidopsis, were the most affected (Figure 1D). We next evaluated the frequencies of nucleotides around the 5’ splice sites (ss) and the 3’ss for all IR events affected in *lsm4-1* compared to the consensus 5 ’ ss or 3 ’ss of all introns in the genome. We found no variation at the 5 ’ ss or 3 ’ss sequences for introns controlled by LSM4, suggesting that LSM4 does not have a prevalent role in splice site recognition (Figure S1A).

When analysing genes differentially expressed (DEGs) in *lsm4-1* compared to wild-type plants, we found 3764 genes (log2 fold change >|1|, FDR <0.05) (Table S3). 25% (238/982) of LSM4 targets had changed splicing patterns, while 15% (121/982) were differentially expressed (Figure 1E), showing for the first time in plants a role of LSM4 in both processes mediated by the two LSM complexes, with a more prominent quantitative effect on splicing. For 33 target genes, binding of LSM4 affects both differential expression and AS (Figure S2B). For example, binding of LSM4 to *AT1G17990* likely stabilises its transcript as loss of *LSM4* causes downregulation of the gene (Figure 1F). At the same time, we observed increased IR of intron 4 in *lsm4-1* (Figure 1F). On the contrary, LSM4 binds to the transcript encoding the MYB59 transcription factor (AT5G59780) which likely destabilised it, as we observed increased levels of all the transcripts isoforms in *lsm4-1* and LSM4 binding causes increased retention of intron 1 (Figure S2C).

### PRMT5-mediated LSM4 methylation associates with a variety of AS events

To address a possible role of R methylation of LSM4 in splicing modulation, we compared splicing alterations in the *prmt5-5* mutant obtained from our recent studies (Mateos *et al*., 2022) (Table S5) and *lsm4-1* mutants analysed here. We reasoned that AS of transcripts in both, *lsm4-1* and *prmt5-5*, could be due to PRMT5-mediated methylation of LSM4. *prmt5-5* shows alteration in 3406 splicing events corresponding to 2601 genes (Mateos *et al*., 2022), with 10% (730 of 6480) of the events affected also in *lsm4-1* (Figure 1G). Interestingly, when we compared the genes in which splicing was affected in both genotypes, the overlap increased to 30% (1348/4499 genes) (Figure 1H). More, differently to what occurs when looking at the whole AS affected in *lsm4-1,* the analysis of the 5’ ss of the 730 shared events disclosed a decrease in the frequency of consensus for the dominant G at the -1 position and a tendency towards randomization of the nucleotides present at the -2 position (Figure S1B), similar to our previous results with the *prmt5-5* mutant (Sanchez *et al*., 2010). It was previously suggested that PRMT5 modulates AS by contributing to the recognition of weak 5’ ss. Interestingly, 50% of LSM4 bound transcripts that have a splicing defect in *prmt5-5* are also affected in *lsm4-1* (Figure 1I). Moreover, the AS affected by *lsm8,* a mutant of a exclusive component of the spliceosomal LSM2-8 complex (Carrasco-López *et al*., 2017), and *lsm4* mutations (this study) cover 65% of the DU genes found in *prmt5*-5 (Figure 1I). Coherently 38% of LSM4 RIP target genes show altered splicing in *lsm4-1* or *lsm8-1* or *prmt5-5*, confirming a strong molecular connection between PRMT5 and the U6 snRNP LSM2-8 complex.

Given the significant overlap in splicing changes between *lsm4-1* and *prmt5-5*, PRMT5 could at least in part affect the function of the U6 snRNP LSM2-8 complex through methylation of LSM4. Noteworthy, our results demonstrate that LSM4 regulates AS by direct binding and in part through PRMT5 action.

### RGG domain mutants of LSM4 rescue the *lsm4-1* phenotype

To reveal to what extent the methylation of Rs in LSM4 is relevant at a molecular and physiological level, we generated transgenic plants carrying a wild-type version of LSM4 or an unmethylable version of LSM4 in the *lsm4-1* background. LSM4 has three arginine-glycine-rich (RGG) motifs containing nine Rs in total at its C-terminus (Figure 2A) which can be methylated by PRMT5 in vitro (Deng *et al*., 2010; Zhang *et al*., 2011). We transformed heterozygous *lsm4-1* mutant plants with wild-type LSM4 (35S::LSM4^R^) and with a version of LSM4 where the nine Rs from the three RGG motifs were changed to lysine (K) (35S:LSM4^RxK^), which maintains the overall charge of the protein but cannot be methylated by PRMT5. Notably, both versions of LSM4, LSM4^R^ and LSM4^RxK^ fully rescued the *lsm4-1* phenotype, generating adult plants that resemble the wild-type plants in normal growth conditions (Figure 2B). To test if changes from Rs to Ks indeed prevent symmetric dimethylation of the LSM4 protein, we performed western blots using protein extracts from LSM4^R^ and LSM^RxK^ seedlings probed with the SYM10 antibody directed against symmetric dimethyl-arginine. Extracts from *prmt5-5* plants that have globally impaired symmetric dimethylation of Rs served as negative control (Mateos et al., 2022). We observed a positive signal at ∼15 kDa in extracts from plants expressing the wild-type LSM4^R^ protein which was absent in *prmt5-5* extracts (Figure 2C). Similarly to *prmt5-5,* LSM4^RxK^ plants showed no methylation signal (Figure 2C), suggesting that mutating Rs to Ks bypasses methylation of LSM4. Still, as no anti-LSM4 antibody is available to verify that the positive band we observed in the western blot is indeed LSM4, and to confirm that the absence of signal in LSM4^RxK^ is not due to altered protein abundance, we generated new plants expressing variants of YFP:LSM4. Similarly to plants used for our RIP-seq experiment, we inserted the LSM4^RxK^ version downstream of the YFP tag. Western blots using 35S::YFP:LSM4^R^ and 35S::YFP:LSM4^RxK^ protein extracts revealed with SYM10 confirmed the results we observed with the untagged lines, where methylation is prevented by changing Rs to Ks (Figure 2D). The expression level of both versions of LSM4 is similar as detected with the GFP antibody (Figure 2D). We previously observed that *lsm4-1* mutant had a long-period phenotype as the circadian period of expression of clock genes such as *CCA1* and *CCR2* genes was lengthened by more than 2 h in *lsm4-1* homozygous mutants relative to wild-type plant (Perez-Santangelo *et al*., 2014). This highlights the role of LSM4 in maintaining the correct function of the clock. We next analysed if methylation of LSM4 is needed to ensure proper clock function. Given that the *lsm4-1* mutant has a small size of its leaves, which does not allow for the measurement of rhythms in leaf movement (Perez-Santangelo *et al*., 2014) we analysed both versions of LSM4, LSM4^R^ and LSM4^RxK^ in the *lsm4-1* background compared to wild-type and found that neither of them showed the distinct long-period phenotype (Figure 2E), suggesting that the methylation status of LSM4 does not impact the role of LSM4 on circadian rhythm. Taken together, these results demonstrate that the Rs of all three RGG domains of LSM4 are methylated in vivo and that changing these residues to Ks prevents methylation without affecting protein stability. More importantly, LSM4 methylation is not critical for rescuing the phenotype under normal growth conditions.

**FIGURE 2.**
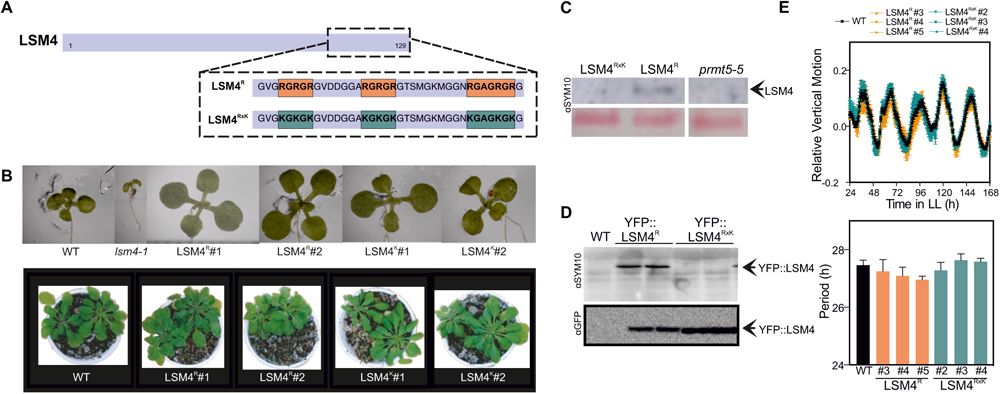
The RGG domains of LSM4 are not required for *lsm4-1* phenotype rescue. (A) Scheme showing wild-type (LSM4^R^) and mutant (LSM4^RxK^) LSM4 C-terminal RGG domains (boxes). (B) Representative pictures of wild-type, *lsm4-1* and two independent lines for each type of transgenic plants, LSM4^R^ and LSM4^RxK^. 12 days-old seedlings (top panel) and adult plants from 4 weeks (bottom panel). (C) Methylation of LSM4 protein is impaired by R to K changes. Immunoblot of LSM4^R^, LSM4^RxK^ and *prmt5-5* samples with the SYM10 antibody. The *prmt5-5* mutant was used as a negative control for impaired methylation. Ponceau red staining shows equal loading across samples. The arrow represents the expected size for LSM4. (D) Immunoblot of wild-type (WT), YFP::LSM4^R^ and YFP::LSM4^RxK^ samples developed with SYM10 (top) or anti-GFP antibody (bottom). The arrow represents the expected size for the YFP:LSM4 fusion protein. (E) Circadian rhythm of leaf movement. Plants’ vertical leaf motion (relative vertical motion: RLM) was obtained for the first pair of leaves of seedlings entrained under long-day conditions (16 h light/8 h dark) and then transferred to constant light (LL). WT, wild-type, (black squares), LSM4^R^, three independent lines from *lsm4-1* mutant plants transformed with LSM4^R^ (orange triangles and diamond) and LSM4^RxK^ , three independent lines from *lsm4-1* mutant plants transformed with LSM4^RxK^ (green triangles and circle) (top panel). The period length of leaf movement rhythms is estimated by Fast Fourier transform–nonlinear least-square test (FFT-NLLS). Error bars represent SEM (bottom panel).

### LSM4 methylation impacts AS of genes associated with stress response

The large overlap of genes with splicing changes between *prmt5-5* and *lsm4-1* mutants (Representation Factor 3.9, p< 6.915e-233) suggests that the activity of LSM4 in controlling a subset of splicing events might be mediated in part by its methylation in the RGG domains (Figure 1). To directly address this, we performed RNA-seq experiments of the transgenic lines expressing the methylable LSM4^R^ or the unmethylable LSM4^RxK^, in the *lsm4-1* background. To analyse our data, we performed three different pairwise comparisons: LSM4^R^ in *lsm4-1* vs *lsm4-1;* LSM4^RxK^ in *lsm4-1* vs *lsm4-1* and *lsm4-1* vs wild-type. More than 90% of the DEGs in *lsm4-1* overlapped with those of either LSM4^R^ or LSM4^RxK^ lines, highlighting that the transcriptomes of LSM4^R^ and LSM4^RxK^ are much alike (Figure 3A, Table S2, S5, S6), which also correlates with the ability of both versions of LSM4 to rescue the *lsm4-1* phenotype under normal growth conditions. Most importantly, when we focused on the impact of the LSM4 methylation status on AS, we found that of the 6480 events altered in *lsm4-1* relative to wild-type (Table S1), 4888 were shared with either LSM4^R^ or LSM4^RxK^ when comparing with *lsm4-1*, reaching 75% of the total events (Figure 3A, Table S7, S8) showing that methylation of LSM4 has a larger effect on splicing than on transcripts levels.

**FIGURE 3.**
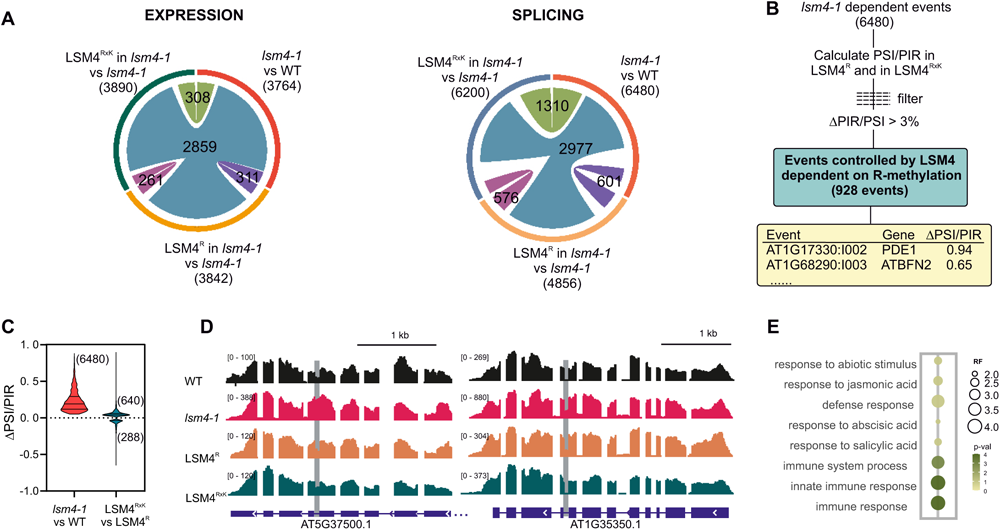
LSM4 R Methylation affects a subset of transcripts involved in stress responses. (A) Summary of the transcriptome analysis is shown as a Circos plot. The total number of differentially expressed genes (left) or affected splicing events (right) in each mutant is shown below the mutant name. Connecting lines are scaled and represent shared affected events. Blue lines shared across all genotypes; green lines shared between LSM4^RxK^ and *lsm4-1*; light violet, shared between LSM4^RxK^ and LSM4^R^ and dark violet shared between LSM4^R^ and *lsm4-1*. Numbers above the lines represent the number of genes (left) or events (right). (B) Scheme showing the rationale behind analysis to define splicing events dependent on LSM4 methylation. (C) The effect of *lsm4-1* mutation is larger than te effect of its methylation. Δ PSI/PIR values for the spliing events affected by LSM4 methylation in *lsm4-1* vs wild-type or LSM4^R^ vs LSM^RxK^. Numbers describe the amount of events. (D) IGV view of mapped reads for selected transcripts affected by R methylation in WT, *lsm4-1,* LSM4^R^ and LSM4^RxK^. The grey line denotes the splicing defect as quantified by RNA-seq analysis. (E) GO-term enrichment analysis of genes affected by R methylation represented as a bubble plot. The size of the bubble is the representation factor (RF).

To examine more precisely the role of LSM4 methylation in splicing, we defined events that depend on LSM4 methylation. For this, we calculated the percentage splice-in, PSI, and percentage intron retention, PIR values for the LSM4^R^ and LSM4^RxK^ genotypes of the 6480 splicing events affected in *lsm4-1* (Figure 3B). Next, we selected events that differ in their PSI or PIR between LSM4^R^ and LSM4^RxK^ by more than 3% (Xin *et al*., 2017). We found 928 splicing events, corresponding to 846 genes, that we called events controlled by LSM4 dependent on R methylation (Figure 3B, Table S9). Of these, we first focus on the 827 IR events, where we observed a significant change in the frequency of the G nucleotide at position -1 at 5’ ss, although to a lesser extent than observed for *prmt5-5* (Figure S1C). Interestingly, when we compared the effect of the *lsm4-1* mutation with that of LSM4 methylation, we found two important characteristics. First, the effect of mutating LSM4 on splicing is larger than that of its methylation, as judged by the higher PSI or PIR values calculated for the same set of shared splicing events (Figure 3C). Secondly, the *lsm4-1* mutation is always detrimental to splicing, meaning that there are more retained introns in the mutant than in the wild-type. Strikingly, LSM4 methylation has a differential effect on IR events. We found that ∼ 70% of the IR events in *lsm4-1* were restored better in LSM4^RxK^ than in LSM4^R^ (Figure 3 C), as shown for *AT5G37500* and *AT1G35350* (Figure 3D), suggesting that methylation could be inhibitory for certain splicing events.

To begin to understand the biological processes that are affected by LSM4 methylation, we focused on AS events with a stronger PSI or PIR difference between LSM4^R^ and LSM4^RxK^ (PSI or PIR difference between genotypes >5%, Table S9, highlighted in blue). We tested this subset of events for overrepresented functional categories by GO term enrichment analysis. Intriguingly, the most significant functional categories were associated with immune responses, including defence-associated salicylic and jasmonic acid hormonal signalling (Figure 3E). In addition, response to drought, salinity and ABA signalling appear as enriched categories, suggesting that LSM4 methylation has a minor effect on plant growth and development, but might be relevant when plants are exposed to adverse conditions (Figure 3E). When we analysed the genes according to the effect on splicing, we observed that the genes from LSM4^RxK^ whose events were more similar to wild-type than the LSM4^R^ to wild-type were almost exclusively enriched in defence-related categories. Collectively, we found that LSM4 methylation plays an important role in modulating gene expression and splicing patterns of a subset of genes associated with biotic and abiotic stress responses, showing a possible role of LSM4 R methylation in the response to these stresses.

### Impaired methylation of LSM4 leads to hypersensitivity to ABA and salt stress

Given that the terms “response to abscisic acid” and “response to abiotic stress” were enriched in our GO terms analysis of genes with splicing events dependent on LSM4 methylation (Figure 3E), we set out to study the effect of LSM4 methylation during abiotic stress. Considering that the increase of ABA levels is a primary signal induced during the adaptive response to abiotic stresses, including salinity (Yang *et al*., 2017) we studied the response of seedlings of the transgenic lines LSM4^R^ and LSM4^RxK^ under ABA and NaCl stress. The addition of ABA did not have any effect on the germination rate after 48 h of treatment among the transgenic lines (Figure S4A), however, LSM4^R^ and LSM4^RxK^ seedlings showed either less or more sensitivity to ABA, respectively, when analysing cotyledon greening, compared to the wild-type plants (Figure 4A-C). Immediately after germination, ABA is associated with a developmental checkpoint inducing growth arrest when young seedlings are stressed (Lopez-Molina *et al*., 2001). LSM4^RxK^ seedlings displayed a strong decrease in fresh weight (FW) after 7 days on 1µM ABA compared to LSM4^R^ and the wild type, confirming the ABA hypersensitivity phenotype of plants impaired in LSM4 methylation (Figure 4D). Similarly, LSM4^RxK^ seedlings when grown on MS plates supplemented with 50 mM NaCl showed a reduction in FW and lateral root (LR) development (Figure 4E-F). No differences between lines were detected at primary root (PR) elongation (Figure S4B). In addition to growth retardation, when plants are exposed to salinity they suffer chlorosis. When treated with 75mM NaCl, LSM4^RxK^ seedlings were more sensitive to salt stress in terms of loss of chlorophyll content compared to LSM4^R^ (Figure 4G).

**FIGURE 4.**
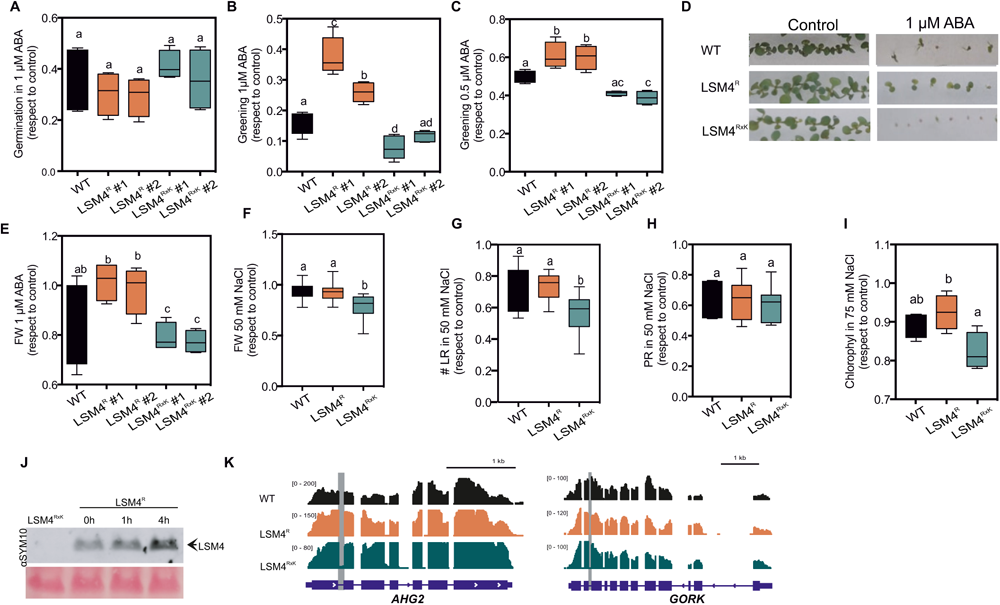
LSM4 R methylation is required for abiotic stress responses. (A-B) Greening of the wild type (WT), two independent transgenic lines of 35S:LSM4^R^ and two independent transgenic lines of 35S:LSM4^RxK^. Seeds were sown in MS supplemented with 1 µM or 0.5 µM ABA as indicated. Greening was analysed after 7 days. (C) Representative seedlings of greening assay at 1µM ABA. (D) Four days-old seedlings were transferred to 1 µM ABA and fresh weight (FW) was quantified 7 days post treatment. (E-F) Four days-old seedlings were transferred to 50 mM NaCl. FW (E) and lateral root (LRs) (F) were analysed 7 days post-treatment. (G) Chlorophyll content of plants treated with 75mM NaCl for 7 days. For all experiments, different letters indicate significant differences among genotypes, p < 0.05 (One-way ANOVA followed by Tukey’s multi-comparison test). (H) Methylation of LSM4 protein is changed upon ABA treatment. Immunoblot of LSM4^R^ samples after 0, 1h or 4h of 10 µM ABA treatment developed with the SYM10 antibody. The LSM4^RxK^ mutant was used as a negative control for impaired methylation. Ponceau red staining shows equal loading across samples. The arrow represents the expected size for LSM4. (I) IGV view of mapped reads for selected transcripts involved in ABA responses affected by R methylation in WT, *lsm4-1,* LSM4^R^ and LSM4^RxK^. The grey line denotes the splicing defect as quantified by RNA-seq analysis.

To further explore the dynamics of LSM4 methylation in vivo, we used the SYM10 antibody to test sDMA levels in protein extracts from LSM4^R^ seedlings incubated in MS medium supplemented with ABA. An induction of LSM4 methylation was detected 4h post-treatment in LSM4^R^ seedlings in vivo (Figure 4H). LSM4^RxK^ was used as a negative control. Previous studies reported that the R methylation signal of some proteins of ∼14kDa was increased in wild-type seedlings subjected to salt and ABA treatments (Zhang et al., 2011, Hu et al., 2017) and that nitrosogluthathione (GSNO) enhanced PRMT5 methyltransferase activity in vitro using histone4 and LSM4 as substrates (Hu *et al*., 2017). Here, our results show that the hypersensitivity to ABA of seedlings impaired in LSM4 methylation correlates with the increase in LSM4 methylation, suggesting that methylation of this U6 snRNP component could play a role in plant responses to abiotic stress mediated by ABA signaling.

### LSM4 methylation differentially controls alternative splicing of specific introns from selected abiotic stress-related pre-mRNAs

Since the unmethylated LSM4^RxK^ lines displayed a different response to ABA and salinity than the methylated LSM4^R^ and WT seedlings, we next explored splicing patterns of genes related to both treatments to determine a possible molecular mechanism behind. Genes selected from GO categories of drought, salinity or ABA response, such as *ABA-HYPERSENSITIVE GERMINATION 2* (*AHG2*) and cell outward potassium channel (*GORK*) showed IR events that are differentially affected in LSM4^R^ compared to LSM4^RxK^ lines, indicating that these IRs are dependent on the methylation status of LSM4 (Figure 4I, Figure S5). *AHG2* is induced by ABA and the corresponding mutant is hypersensitive to ABA in germination assays (Lopez-Molina *et al*., 2001; Hirayama *et al*., 2013), while the mutant for *GORK* gene has an increased water consumption and reduced FW in response to ABA, similar to the phenotypes reported here for LSM4^RxK^ seedlings (Figure 4A-E). Therefore, we propose that impaired splicing of these genes in LSM4^RxK^ lines may contribute to ABA hypersensitivity. Furthermore, we analysed other genes involved in ABA signalling and related to abiotic stress responses, finding that *RESPONSIVE TO DESICCATION 22* (*RD22*) splicing is regulated by PRMT5 but not by LSM4. While splicing of *ABSCISIC ACID RESPONSIVE ELEMENTS-BINDING FACTOR 2* (*ABF2; AT1G45249*) and HYPERSENSITIVE *TO ABA1* (*HAB1; AT1G72770*) are regulated by LSM4 (Figure S6) similar to what was found by Zhang *et al*., 2011, but only *ABF2* seems to be dependent on LSM4 methylation. Taken together, LSM4 methylation regulates intron splicing of genes associated with ABA, such as AHG2, and ABF2, which may contribute to plant adaptation to abiotic stress including salinity and drought.

### Methylation of LSM4 negatively regulates splicing of defence genes and plant immunity

Arabidopsis plants respond to bacterial infection via two connected branches of immune signalling. The pathogen-associated molecular pattern (PAMP)-triggered immunity (PTI) is initiated when plasma membrane-localised pattern recognition receptors detect the corresponding PAMPs; while effector-triggered immunity (ETI) is a stronger response and is activated through recognition of virulence factors by intracellular immune receptors including the nucleotide-binding leucine-rich repeat (TIR-NB-LRR) protein family, and induces specific and localised immune reactions (Boller and Felix, 2009; Nakai *et al*., 2013). Since our GO analysis of genes with methylation-dependent splicing events of LSM4 showed an enrichment of the terms “immune responses”, including defence-associated “salicylic and jasmonic acid hormonal signalling” (Figure 3E), we focused on whether LSM4 methylation has a regulatory role during bacterial infection. We first compared genes differentially spliced dependent upon the LSM4 methylation status (Table S9) with genes differentially spliced in wild-type plants after infection with *Pseudomonas syringae* pv*. tomato* DC30000 (*Pst*) (Golisz *et al*., 2021). For 113 genes splicing is affected by the LSM4 methylation status as well as *Pst* infection (Figure 5A, Table S10), implying that R methylation can modulate bacterial resistance by regulating the splicing of specific transcripts. Interestingly, 87 out of 113 genes had increased levels of the IR isoform in the LSM4^R^ genotype judged by their PIR value (Table S11, Figure 5B). LSM4^R^ had increased IR of *VASCULAR PLANT ONE ZINC FINGER PROTEIN* (*VOZ1*) and *RADICAL-INDUCED CELL DEATH1* (*RCD1*) mRNAs, both required for the activation of the immune responses to bacterial and fungal pathogens in Arabidopsis (Bardou *et al*., 2014; D’Ippolito *et al*., 2016; Wirthmueller *et al*., 2018). In contrast, IR of the *IAA-LEUCINE-RESISTANT (ILR1)-LIKE 3* (*ILL3*) member of the amidohydrolase family responsible for increasing the levels of free active IAA and negatively associated with defence (D’Ippolito *et al*., 2016) is higher in LSM4^RxK^ seedlings compared to LSM4^R^ (Table S11). So far, these results indicate that LSM4 methylation modulates IR events of genes involved in plant immunity by regulating the ratio of isoforms to increase or decrease the functional isoform depending on its activity in plant defence. In agreement, a significant number of the introns differentially targeted by LSM4 methylation were in transcripts encoding defence signalling proteins associated with the ETI branch of the immune response (Figure 3E). In all cases, intron retention in these defence signalling genes was higher in LSM4^R^ than in unmethylable LSM4^RxK^ plants. For instance, five members of the TIR-NBS-LRR family show increased IR in LSM4^R^ compared to wild-type, while LSM4^RxK^ plants show more efficient splicing almost indistinguishable from wild-type plants (Figure 5C). For all of these genes, IR causes a premature stop codon leading to shorter proteins (Figure S7), meaning that in LSM4^R^ plants, intron-containing transcripts would encode premature stop codons that may result in truncated proteins with potentially diminished functions. Furthermore, ZAR1, a key NLR protein that is a calcium-permeable cation channel for triggering immunity and cell death, shows increased IR in plants expressing LSM4^R^. Interestingly, unmethylable LSM4^RxK^ plants show perfect splicing of intron 1, increasing the level of the functional isoform that upregulates plant defences (Figure 5D). These results suggest that LSM4 methylation could be a means for controlling proper splicing of defence genes during bacterial infection. To test this, leaves from two-week-old LSM4^R^ plants were infiltrated with the bacterial pathogen *Pst* or MgSO_4_ as mock. We observed that 24 h post-infection, plants infected with *Pst* displayed a reduction in LSM4 methylation compared to mock-treated plants as evidenced by immunoblots with SYM10 antibody (Figure 5E). An early increment of LSM4 methylation 4 h post-infection, which is opposite to the 24h response, was also detected. Still, this induction could be triggered by bacteria and counteracted by the plant to overcome the infection. This temporal pattern was reported for the accumulation of the phytohormone auxin which is induced by most pathovars of *Pst* (Thilmony *et al*., 2006; Chen *et al*., 2007), while auxin signalling is repressed by Arabidopsis plants to reduce bacterial infection (Navarro *et al*., 2006).

**FIGURE 5.**
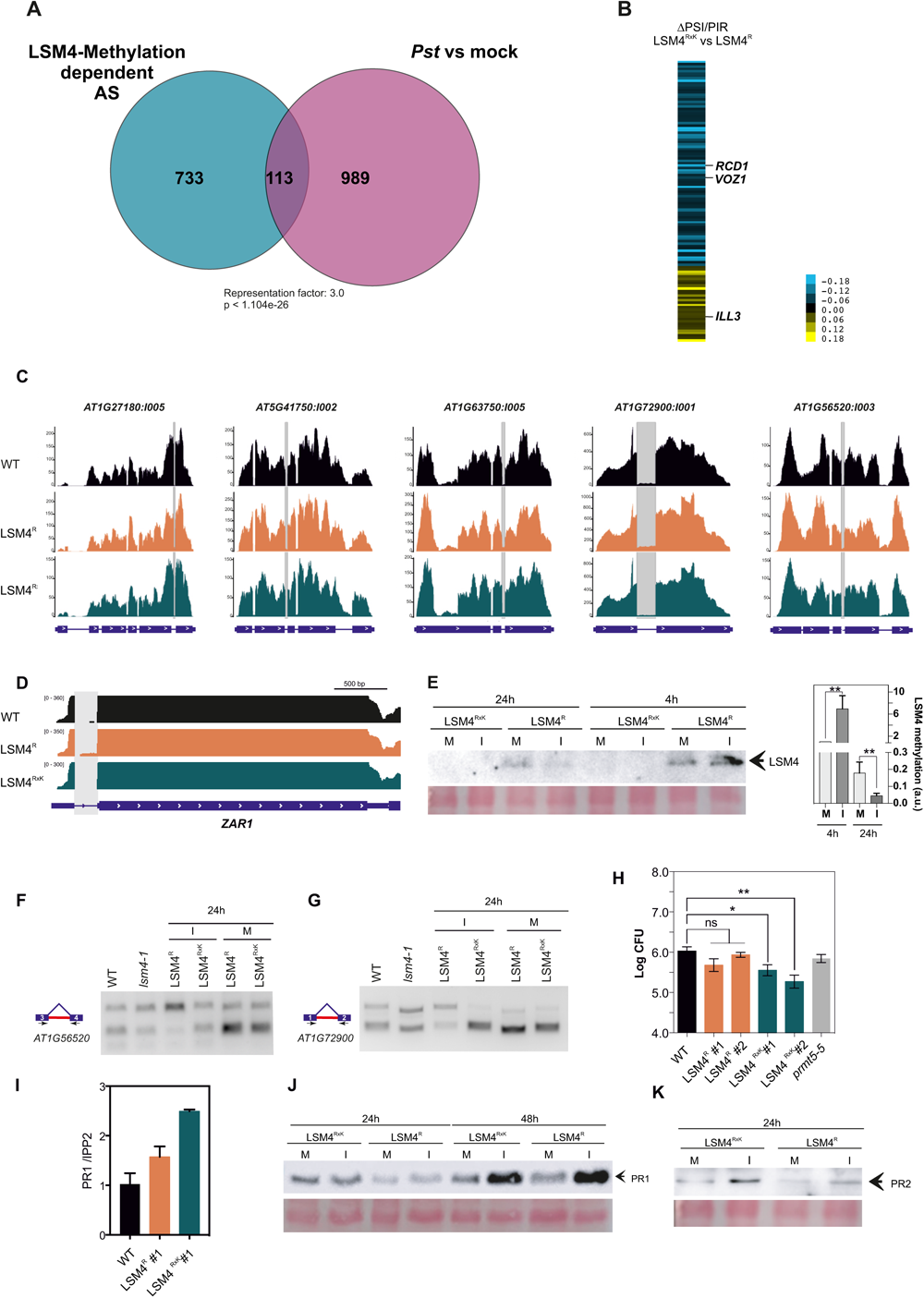
LSM4 methylation modulates plant immunity. (A) Venn diagram showing the extent of overlap for pre-mRNA splicing events affected by LSM4 methylation and those altered upon bacterial infection from Golisz et al, 2021). (B) Heatmap showing ΔPSI/PIR values calculated for the 113 genes from (A) in LSM4^RxK^ vs LSM4^R^. (C) IGV view of mapped reads for selected transcripts coding for TIR-NBS-LRR proteins in WT, *lsm4-1,* LSM4^R^ and LSM4^RxK^. The grey line denotes the splicing defect as quantified by RNA-seq analysis. (D) IGV view of mapped reads for *ZAR1* in WT, *lsm4-1,* LSM4^R^ and LSM4^RxK^. The grey line denotes the splicing defect as quantified by RNA-seq analysis. (E) (Left) Methylation of LSM4 protein upon 4 and 24h of bacterial infection with *Pseudomonas syringae* DC3000. Immunoblot of LSM4^R^ samples with the SYM10 antibody. The LSM4^RxK^ mutant was used as a negative control for impaired methylation. Ponceau red staining shows equal loading across samples. Arrow represents expected size for LSM4. (Right) Quantitation of three independent experiments (** p<0.001; t-Test to mock). (F, G) RT-PCR to detect splicing defects for *AT1G56520* and *AT1G72900* in LSM4^R^ and LSM4^RxK^ upon bacterial infection or mock treatment. The wild-type and *lsm4-1* genotypes were measured under controlled conditions to show IR is increased in *lsm4-1.* Alternative regions are highlighted in red in the diagrams next to the gels and position of the primers used are depicted. (H) Four weeks-old plants grown in long-days conditions were infected by infiltration with *P. syringae* DC3000. Bacterial growth was assessed 2 d.p.i. (CFU: colony-forming units) in WT, two independent lines of LSM4^R^, two independent lines LSM4^RxK^ plants. Data represent the average of log-transformed bacterial growth (n = 8 independent biological replicates). This experiment was repeated twice with similar results. Error bars indicate SEM. p < 0.05 (One-way ANOVA followed by Tukey’s multi-comparison test). (I) Expression levels of *PR1* measured by qRT-PCR. The analysis was conducted in WT, LSM4^R^ and LSM4^RxK^ plants grown under a long-days photoperiod 48-hours post-infiltration with a mock solution (10 mM MgSO4) (n= 3 biological replicates). Results were calculated relative to *IPP2* transcript levels. Levels of PR1 (J) and PR2 (K) proteins upon bacterial infection determined by western blots. PR1 levels were determined in LSM4^R^ and LSM4^RxK^ plants after 24 h or 48 h upon bacterial infection. Levels of PR2 were measured after 24h of infection. anti-PR1 and anti-PR2 antibodies were used respectively. Ponceau red staining shows equal loading across samples.

To better understand the potential molecular consequences on splicing of reduced methylation after infection, we next tested splicing patterns upon 24 h of bacterial infection in two genes of the TIR-NBS-LRR family (*AT1G56520* and *AT1G72900,* Figure 5C). In normal conditions, *lsm4-1* mutant presents an increased IR on *AT1G56520* transcript, which can be visualised by RT-PCR with primers binding to exon 3 and 4 expanding intron 3, compared to wild-type plants (Figure 5F). Upon *Pst* infection, IR increased in LSM4^R^ while LSM4^RxK^ showed wild-type levels of spliced intron (Figure 5F). Similar results were observed when analysing IR in intron 1 of *AT1G72900* gene, which is again increased in LSM4^R^ compared with LSM4^RxK^ under *Pst* infection (Figure 5G). These differences in splicing patterns are given by the methylation status of LSM4 and not by variance in expression of the transgenes, as the *LSM4* transcript accumulates to similar levels in the LSM4^R^ and LSM4^RxK^ lines analysed here (Figure S8). The function of the rest of the genes in plant immunity and the possible effect of IR on its function remain to be analysed.

We reasoned that if increased methylation of LSM4 is detrimental to proper splicing of plant defence genes on one hand (Figure 5F-G) and plant responses to bacterial infection tend to minimise LSM4 methylation in part by reduced levels of PRMT5 on the other hand (Hong *et al*.,2010), this should translate into a differential activation of immune responses in LSM4^R^ and LSM4^RxK^ plants. To assess this, LSM4^R^ and LSM4^RxK^ plants were challenged with the pathogen *Pst*. Leaves were infiltrated with *Pst* and bacterial growth was analysed two days post-infection. We found that unmethylated LSM4^RxK^ lines developed less susceptibility to *Pst* infection compared to LSM4^R^ or wild-type plants (Figure 5H), antagonistically to the abiotic stress response where LSM4^R^ seedlings show better performance compared to unmethylated LSM4^RxK^ seedlings. The enhanced resistance of the LSM4^RxK^ plants was associated with an increased basal expression of the typical marker of the salicylic acid-mediated defence system, *PATHOGENESIS-RELATED 1* (*PR1*) at mRNA and protein levels (Figure 5I-J). In addition, an early response of PATHOGENESIS-RELATED PROTEIN 2 (PR2) was detected at 24 hpi in LSM4^RxK^ infected plants compared to LSM4^R^ (Figure 5K). Wang et al. (2021), found that plants overexpressing LSM4 were more susceptible to bacterial infection which correlated with a reduced expression of *PR1*, *PR2* and *PR5*. Although these results are in line with our observations, we could not detect an increased susceptibility of our LSM4^R^ plants to *Pst* infection compared to the wild-type, probably due to differences in growth conditions. Notably, both results indicate that higher LSM4 levels or increased methylation status seem to downregulate plant immunity and PR activation. This aligns with the effect of R methylation on AGO2 by PRMT5 where plants lacking the RGG domain of AGO2 are more resistant to bacterial infection (Hu *et al*., 2019). Indeed, similar to LSM4^RxK^ plants, *prmt5-5* mutants showed a weak but enhanced response to bacterial infection (Hu *et al*., 2019). This downregulation of PRMT5 during infection could entail reduced LSM4 methylation. Our results show that the plant response to biotic stress could be mediated by splicing regulation of defence response genes through modulation of the LSM4 methylation status.

## DISCUSSION

Alternative pre-mRNA splicing is an important mechanism to rapidly adjust the transcriptome to environmental changes, especially allowing rapid adaptation to stresses without the obligation/need of de novo transcription. R methylation of spliceosomal RBPs is a well-documented post-translational modification (Deng *et al*., 2010; Hu *et al*., 2019; Cao *et al*., 2022). Yet, experimental evidence showing its impact on splicing was still lacking. Using LSM4, an R-methylable U6 snRNP component of the spliceosome as a paradigm for studying RBP methylation, allowed us to uncover an important regulatory layer involved in fine-tuning plant responses to abiotic and biotic stress in an opposite way.

Using RIP-seq, we identified 952 target transcripts of LSM4, many of which are coding for splicing related proteins or RBPs. Cross-referencing the targets with the transcriptome of the *lsm4-1* mutant revealed that for ∼25% of the LSM4 targets splicing was altered, whereas for ∼15% expression levels were changed. Thus, we show for the first time that LSM4 is involved in both AS and steady-state levels of transcripts by directly binding to RNAs, a distinguishing feature that was only described for a small number of RBPs in plants (Staiger *et al*., 2003; Day *et al*., 2012; Bardou *et al*., 2014; Wu *et al*., 2016). The glycine-rich domain of LSM4 was shown to enhance mRNA stability and alter mRNA decay pathway in *Saccharomyces cerevisiae*, but its action is not by binding to the transcripts but by association with the decapping activator Edc3 (Huch et al, 2016).

A close analysis of the LSM4 targets suggests that LSM4 regulates AS and/or mRNA levels of several other RBPs involved in posttranscriptional processes, suggesting a feedback/cooperation among RBPs complexes to ensure a proper transcriptome. Of the LSM4 target transcripts that are as well differentially expressed in the mutant, a similar proportion was up-or down-regulated (Figure S8), suggesting that LSM4 binding can lead to both stabilisation or destabilisation of transcripts without any preference. This does not seem to be a general rule for other RBPs involved in splicing such as *At*GRP7, as in this case, binding reveals a predominantly negative effect of *At*GRP7 on its targets (Meyer et al, 2017).

A possible role of R methylation of LSM4 by PRMT5 in modulating splicing should be reflected by AS changes occurring in both, *lsm4-1* and *prmt5-5*. We found that around 30% of genes differentially spliced in *lsm4-1* are also differentially spliced in *prmt5-5.* Thus, PRMT5 could at least partially affect the function of the U6 snRNP LSM2-8 complex through LSM4 methylation. Methylation of other LSM components of the complex seems less likely. For instance, we could not detect sDMAs on LSM8 (Figure S9). Previously, LSM4, LSM7, LSM6B and LSM8 were detected by mass spectrometry of proteins immunoprecipitated with sDMA-specific SYM11 antibody (Boisvert et al., 2003), but methylation by PRMT5 in vitro was only confirmed for LSM4 (Zhang et al., 2011). Therefore, Zhang et al. (2011) proposed that the unmethylated LSM proteins co-immunoprecipitated non-specifically with methylated LSM4.

LSM4 methylation by PRMT5 may not be completely essential for the subset of splicing events regulated by LSM4 through the U6 snRNP complex since many genes are spliced correctly independently of PRMT5 action (Figure 1G-H). Rather, sDMAs on LSM4 could constitute an extra layer of control that contributes to splicing of specific transcripts, particularly those with weak 5’ss. It should not be discarded that some of the events affected in *lsm4-1* might be an indirect action caused by either altered gene expression, as LSM4 is as well part of the LSM1-7 complex controlling mRNA stability, or by variations on splicing pattern of splicing-related factors due to the *lsm4-1* mutation. In particular, R methylation of LSM4 by PRMT5 does not appear to be a requirement for splicing modulation by the U6 snRNP complex under normal growth conditions. On the contrary, we show that treatments associated with stress are capable of modulating LSM4 methylation levels precisely. LSM4 methylation is beneficial during salinity (Figure 4) while responses to bacterial infection are favoured when methylation levels are limited (Figure 5). This is consistent with the opposite performance of *prmt5-5* and *lsm4-1* mutants under the same stresses (Zhang et al., 2011). Our work suggests that increased R methylation is advantageous for plants under ABA or NaCl, both treatments typically associated with abiotic stresses. Several pieces of evidence support our idea. First, LSM4 sDMA is favoured upon ABA treatments (Figure 4H). Second, unmethylable LSM4 plants show hypersensitivity when facing abiotic stress (Figure 4A-G, Figure S3). Last but most importantly, we found that LSM4 methylation regulates intron splicing of several genes associated with ABA such as *AHG2* and *ABF2*. In abiotic stress IR is increased in LSM4^RxK^ compared to LSM4^R^ or wild-type, indicating that improper splicing of these genes might be responsible for the hypersensitive phenotype (Figure 4K). Furthermore, analysis of the *lsm8* mutant revealed that the U6 snRNP LSM2-8 complex is necessary for the precise splicing of a set of stress-related pre-mRNAs, related to cold and salt stress tolerance (Carrasco-López et al., 2017). Moreover, mutants in *PRMT5*, *LSM4* and *LSM5* are hypersensitive to salinity and ABA due to inaccurate splicing of stress-related genes (Zhang et al., 2011; Cui et al., 2014).

Our results are also in agreement with multiple pieces of evidence associating different components of the spliceosome with redundant and complex regulation of splicing under abiotic stress. The Sm core protein SmEb is induced by ABA treatment, and also modulates the AS of *HAB1* similar to the *SKIP* splicing factor (Huertas et al., 2019; Hong et al 2021; Zhang et al 2022). In addition, an analysis of thousands of public RNA-seq libraries selected *PRMT5* and *SKIP* as the main regulators of introns present in nuclear-unspliced chromatin-bound polyadenylated transcripts, called post-transcriptionally spliced introns (pts) (Jia et al., 2020). Regulation of pts introns by *PRMT5* and *SKIP* could constitute a rapid mechanism to produce fully spliced functional mRNAs for plants to rapidly adapt to changing environmental conditions where pts introns are enriched. We propose that this action of PRMT5 could be in part due to its role as methyltransferase of RBPs.

Contrarily, biotic stress negatively regulates LSM4 methylation, evidenced by reduced SYM10 signal of LSM4 upon bacterial infection (Figure 5E). Consequently, plants expressing unmethylable LSM4 show stronger resistance to *Pst* (Figure 5). A recent study also demonstrates that overexpression of LSM4 renders plants more susceptible (Wang *et al*., 2021), in line with the idea that increased LSM4 methylation is detrimental for plant immunity. All this is in accordance with previous observations that *PRMT5* transcript levels are slightly reduced during infection with *Pst* (Hu et al, 2019). In fact, R methylation of ARGONAUTE2 protein by PRMT5 has been proposed to participate in defence responses. Down-regulation of *PRMT5* during infection causes reduced AGO2 R methylation, leading to stabilisation of AGO2 and AGO2-associated sRNAs to promote plant immunity (Hu et al, 2019). Along with decreased LSM4 methylation levels, we observed that splicing of defence-related genes is also influenced by the methylation status (Figure 5). In general, we found IR of genes involved in plant immunity to be altered following the LSM4 methylation pattern. Interestingly, the IR ratio between isoforms does not always shuffle in the same direction for all genes. For instance, negative regulators of immunity have more IR, while positive regulators have less IR in unmethylable plants or after pathogen entry. But still how and whether an intron is selected for increased or diminished IR upon LSM4 methylation is a question yet to be answered.

Taken together, this is the first direct evidence that R-methylation of a component of U6 snRNP forms an additional layer in post-translational regulation, with antagonistic roles in the plant response to abiotic and biotic stress, respectively (Figure 6). Furthermore, we correlated LSM4 methylation with splicing changes. Thus, PRMT5-mediated methylation of LSM4 has an impact on several stress responses associated with the splicing control of genes involved in these stresses.

**FIGURE 6.**
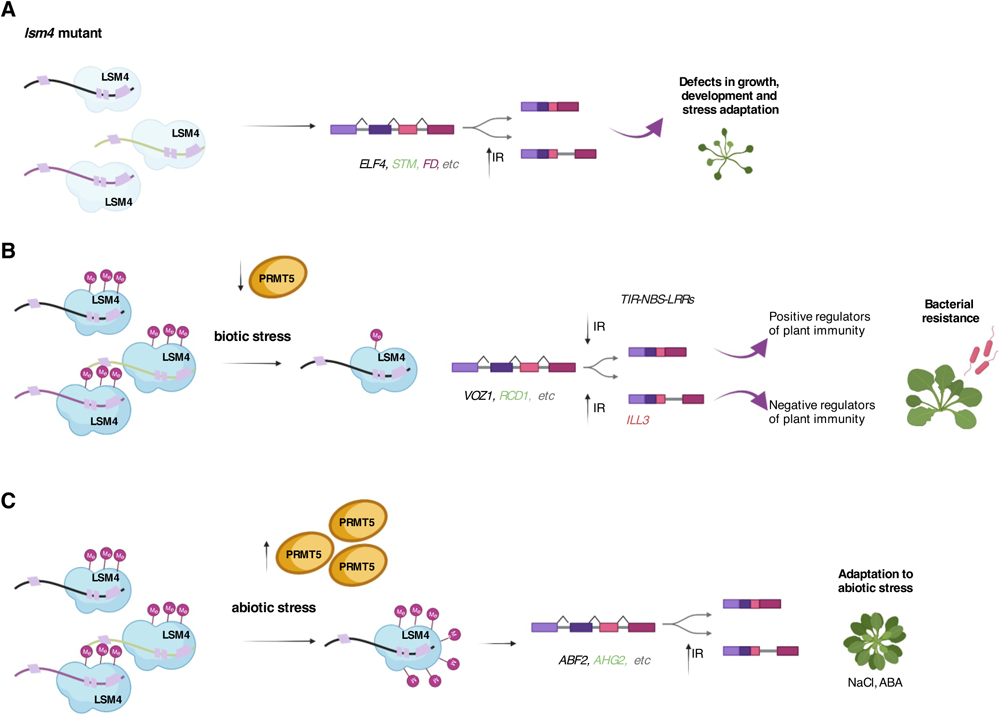
Proposed model for the role of LSM4 methylation in modulation of stress responses. (A) In the absence of *lsm4* a wide range of AS defects leads to aberrant phenotypes and altered stress responses. Splicing defects are enriched in IR events. (B) Upon exposure to bacterial infection, *PRMT5* decreases (Huang et al, 2019) and methylation of LSM4 is reduced (this work). This reduction affects splicing of genes involved in plant immunity to increase bacterial resistance. (C) Treatments associated with abiotic stress increase LSM4 methylation (this work and Hu et al, 2019). A decrease in IR of genes involved in abiotic stress responses leads to an increase of the functional isoforms correlating with improved adaptation to abiotic stress.

## Materials and methods

### Plant material and growth conditions

All *Arabidopsis thaliana* mutants are in the Columbia-0 (Col-0) ecotype. The *lsm4-1* (SALK_063398) and *prmt5-5* mutants were previously described (Perez-Santangelo *et al*., 2014; Sanchez *et al*., 2010). Plants were grown on the soil at 22°C under long days (LD cycles (16 h light/8 h dark); 80 μmol.m^−2^s^−1^ of white light) for infection assays. For the rest of the experiments, seeds were surface sterilised, sown in Murashige and Skoog (MS) medium containing 0.8% (w/v) agar and stratified for 3 d in darkness at 4°C. Seedlings were grown in a growth chamber under controlled conditions

### Generation of transgenic lines

The sequences for LSM4^R^ and LSM4^RxK^ were synthesized and cloned into the pDONR/Zeo (Invitrogen) via the Gateway method. The LSM4^RxK^ sequence was generated by changing the R codons to K codons in the 9 methylable RGG domains. The 35S: LSM4 expression vectors were generated by introducing the AtLSM4^R^ or AtLSM4^RxK^ sequence into the binary vector pEarleyGate 100 by LR recombination. This plasmid was then introduced into *Agrobacterium tumefaciens* GV3101 strain and transformed into *lsm4-1* heterozygous plants using the floral dip method (Clough and Bent, 1998). Transgenic lines were selected on MS media supplemented with 15 mg/L phosphinotricin (Duchefa) and genotyped for the *lsm4-1* mutation using primers described in Table S11.

### Circadian Leaf Movement Analysis

For leaf movement analysis, seeds were sown into small circular pots (filled with growth media) and entrained in LD (16 h light/8 h dark) conditions at 22^◦^C for seven days until the first pair of leaves were fully expanded. Then, plants were transferred to continuous light (LL) and constant temperature (22^◦^C) conditions. Photos were taken every hour for six days and analysed by recording the plants’ vertical leaf motion (relative vertical motion: RLM) with a program developed in java (Iserte and Yanovsky, 2017) based on the Matlab code from the software TRiP: Tracking Rhythms in Plants (https://github.com/KTgreenham/TRiP) (Greenham *et al*., 2015) The circadian period was obtained via fast Fourier transform nonlinear least-squares (FFT-NLLS) analysis using the online program BioDare2 (www.biodare2.ed.ac.uk) (Zielinski *et al*., 2014) The first 24 h were excluded from the analysis to remove potential noise caused by the transfer from entrainment conditions to constant conditions.

### RNA extraction and quantitative real-time PCR

Total RNA was isolated from seedlings using TRIZOL (Ambion) and treated with RNAse-free DNase I (Promega) to remove residual genomic DNA. One microgram of total RNA was used for reverse transcription (RT; Superscript II, Invitrogen). Transcript levels were quantified by quantitative PCR in a Stratagene MX3005P instrument (Agilent Technologies) using *PP2A* (*AT1G13320*) or *IPP2* (*AT3G02780*) as the housekeeping gene. The sequences of the primers used to quantify the expression are listed in Table S11.

### Validation of splicing events

cDNAs were synthesised as above but with SuperScript (SS) III (Thermofischer). PCR amplification was performed using SuperFidelity Taq Enzyme (Invitrogen). Primers used for amplification are detailed in Table S11. RT-PCR products were incubated with SYBR Green before electrophoresis on 2.5% (w/v) agarose gels. We selected the events to be validated according to the following parameters: the gene must have at least 50 reads to ensure good expression, the difference between the isoforms must not be greater than 10% in size, and when possible the neighbouring intron, with no changes, must be included in the PCR product to show that the measured event is specific.

### RNAseq analysis

For RNA-seq experiments, wild-type, *lsm4-1,* LSM4^R^ *and* LSM4^RxK^ in *lsm4-1* background seedlings were grown on MS medium containing 0.8% (w/v) agar for 12 days under continuous light (LL) at 22°C. Three biological replicates were collected, whole seedlings were harvested and total RNA was extracted with RNeasy Plant Mini Kit (QIAGEN) following the manufacturer’s protocols. To estimate RNA concentration NanoDrop 2000c (Thermo Scientific) was used. Libraries were prepared and sequenced at the Max Planck-Genome-Centre Cologne (MP-GC).

### RIP-seq analysis

RIP was done as previously described except for the bioinformatic analysis (Meyer *et al*., 2017). We calculated the enrichment of the signal from IP using 35S:YFP:LSM4 plants over an IP from the negative control plants 35S:YFP. Briefly, plants grown in MS plates in LL for 12 days were vacuum-infiltrated with 1% formaldehyde for 15 min followed by quenching with 125 mM glycine. A whole-cell extract was prepared in RIP-lysis buffer. The extract was pre-cleared with Sepharose beads and subjected to immunoprecipitation with GFP-Trap beads (Chromotek). After extensive washing with RIP washing buffer, co-precipitated RNAs were extracted with Trizol (Invitrogen) and treated with DNase (Promega). Libraries were prepared and sequenced at the Max Planck-Genome-Centre Cologne (MP-GC).

### Alternative splicing and differential expression analysis

Alternative splicing (AS) analysis was performed as previously described in Mateos *et al*., 2022 using R with the ASpli package (Mancini *et al*., 2021). For expression analysis, genes with a false-discovery rate (FDR) <0.05 and log_2_FC >|1.0|) were considered as differentially expressed (DEGs) between genotypes. For splicing analysis, PSI (percent of inclusion) and PIR (percent of IR) were calculated and differential splicing was considered for bins with FDR <0.15 and ΔPSI/PIR >0.05. For the analyses of RNA-seq data from *lsm8*, bam files from our previous work (Carrasco-López *et al*., 2017) were treated as described above to compare splicing events.

### Physiological response to ABA and salinity

Seeds were sown on MS medium supplemented with 0.5 or 1 µM ABA. The proportion of germinated seeds was scored after 48 h while greening after 7 d of treatment. Both parameters correspond to radicle emergence and fully opened cotyledons, 0.5 and 1 stages respectively, according to Boyes et al (2001). For the analysis of primary root (PR) elongation, lateral root (LR) formation and fresh weight (FW), 4-d-old seedlings were grown vertically on MS agar plates and then transferred to MS supplemented with indicated concentrations of ABA or NaCl. Quantifications were assayed after 7 d. Chlorophyll content was measured spectrophotometrically at 645 and 663 nm according to Arnon (1949).

### Infection assay

Five-week-old plants were inoculated into the abaxial side of 8 to 10th leaves with a bacterial suspension of *P. syringae* pv. *tomato* DC3000 (OD600 = 0.0002) or mock solution (10 mM MgSO_4_). Three disks were punched from each leaf 48 h post-infection (h.p.i) and bacterial growth was determined by counting bacterial colonies in plate assays according to de Leone et al (2020).

### Methylation status and PRs level

Seven days-old seedlings were transferred to liquid MS supplemented with 10 µM ABA for indicated times and assayed for LSM4 methylation status by anti-SYM10 immunoblot. Five-week-old plants were inoculated with a bacterial suspension of *P. syringae* pv. *tomato* DC3000 (OD600 = 0.002) or mock solution (10 mM MgSO4), harvested 4, 24 and 48 h.p.i. and assayed for LSM4 methylation status and PR induction by immunoblots. Total proteins were extracted in extraction buffer (50 mM Tris-HCl pH 7.2, 100 mM NaCl, 10% (v/v) glycerol, 0.1% (v/v) Tween-20, 1 mM phenylmethylsulfonyl fluoride (PMSF) and plant protease inhibitor (Sigma)) and then centrifuged at 12,000 g and 4°C for 15 min. Equal amounts of protein were loaded onto SDS-PAGE, electro-transferred onto nitrocellulose membranes and probed with anti-SYM10 (Millipore, 07-412), anti-PR1 (Agrisera, AS10 687) or anti-PR2 (Agrisera, AS12 2366) antibodies overnight followed by secondary anti-rabbit antibody coupled to peroxidase (Invitrogen, USA). Proteins were visualised using the ECL kit (Pierce, USA) in an ImageQuant LAS 4000, GE Healthcare. Ponceau staining is shown as loading control.

### IP and methylation status of LSM4-GFP and LSM8-GFP fusion proteins

Protein was isolated from 2-week-old *35S:LSM4-GFP* or *35S:LSM8-GFP* transgenic lines in IP buffer containing 50 mM Tris-HCl, pH 7.5, 150 mM NaCl, 10% glycerol, 0.1% Nonidet™ P40; 1 mM phenylmethylsulfonyl fluoride (PMSF), and protease inhibitor cocktail (Sigma-Aldrich, USA). After centrifugation at 12,000 g and for 15 min at 4°C, protein extracts were incubated with GFP-Trap Agarose (Chromotek, Germany) at 4°C for 1 h with gentle rotation. After centrifugation at 2500 g for 4 min, an aliquot of the supernatant was saved for flow-through control. Beads were washed 4 times in IP buffer and bound proteins were extracted in 2.5X Laemmi SDS-sample buffer. R-methylated LSM proteins were identified by immunoblot with anti-SYM10 as previously described above.

## Data availability

The RIP-seq and RNA-seq raw data have been deposited in ArrayExpress (Kolesnikov et al., 2015) at EMBL-EBI (www.ebi.ac.uk/arrayexpress), under accession numbers E-MTAB-12369 (RIP-seq dataset) and E-MTAB-12370 (RNA-seq dataset).

## Acknowledgements

We thank Dr. Marlene Reichel for critical reading of the manuscript. We thank Kristina Neudorf for technical assistance on RIP and Martin Lewinski for bioinformatic assistance.

## Author contributions

JLM, MJI, DS and MJY designed the research; YCA, MJI, SPS, MdeL, RC, TK, and JLM performed the experiments, JLM performed the bioinformatic analysis, JLM, MJI and SPS wrote the paper.

## SUPPLEMENTARY FIGURES

**Figure S1. Bioinformatic analysis of 3’and 5 splice-site sequences.**

(A) Pictograms showing the frequency distribution of nucleotides at the 5′ splice site of 119,072 GT_AG_U2 Arabidopsis introns and 3’splice site of 173394 AG sites (top) or the most significantly intron retention events whose splicing were altered in *lsm4-1* (bottom). (B) Pictograms showing the most significantly intron retention events altered in *lsm4-1* and *prmt5-5.* (C) Pictograms showing the most significantly intron retention events affected by LSM4 methylation. The representation factor (RF) is the frequency in the data set of interest divided by the total frequency. For each RF a p-value was calculated using the hypergeometric test.

**Figure S2. Direct binding of LSM4 to transcripts alters both AS and mRNA levels.**

(A) List of genes bound by LSM4 that are affected in AS and mRNA levels simultaneously. (B). IGV browser showing read density at the *AT5G5980* locus for RIP-Seq and RNA-Seq experiments. This constitutes an example of an LSM4 target with impact on splicing and mRNA steady-state levels.

**Figure S3. LSM4 methylation under abiotic stress**

(A) Seed germination of the wild type (WT), two independent transgenic lines of 35S:LSM4^R^ and two independent transgenic lines of 35S:LSM4^RxK^. Seeds were sown in MS supplemented with 1 µM ABA. (B) Four days-old seedlings were transferred to 50 mM NaCl and primary root (PR) growth was measured 48h post treatment. For all experiments, different letters indicate significant differences among genotypes, p < 0.05 (One-way ANOVA followed by Tukey’s multi-comparison test).

**Figure S4. LSM4 affects splicing of abiotic stress related genes.**

Integrated genome browser view showing coverage plots from selected genes related to abiotic stress with splicing defects in *lsm4-1.* Data were obtained in this study from seedlings grown under continuous light.

**Figure S5. Effect of LSM4, PRMT5 and LSM4 methylation on splicing of abiotic stress related genes.**

(A-C) Integrated genome browser view showing coverage plots from three selected genes with splicing defects in *prmt5-5* (A), *lsm4-1* (B) or by LSM4 R methylation (C). Data for *prmt5-5* mutants were obtained from (Mateos et al, 2022). Data for the LSM4 related genotypes were obtained in this study. The alternatively spliced intron is highlighted with a box.

**Figure S6. Representative models and retained intron splice forms of the five TIR-NBS-LRR genes.**

Exons are shown as boxes and introns are designated as lines. Intron retention events are highlighted in cyan. The effect of intron retention is depicted below the representative model. STOP codons are indicated. Bordeaux areas indicate the coding region and grey areas UTRs.

**Figure S7. *LSM4* transcript levels in transgenic lines.**

Accumulation of *LSM4* transcripts in 10-d-old wild-type, LSM4^R^, LSM4^RxK^ and *lsm4-1* seedlings grown under control conditions at 22°C. Data represent mean±SEM.

**Figure S8. *LSM4* transcript levels in transgenic lines.**

Expression changes of LSM4 direct targets in *lsm4-1* mutant.

**Figure S9. LSM8-GFP is not methylated at its arginines.**

Immunoprecipitation (IP) assays in fluorescent protein-tagged LSM4 and LSM8 overexpressing lines with anti-GFP antibody. Wild-type plants were used as negative controls. Immunoblots with anti-GFP and anti-dimethyl arginine antibody, symmetric, SYM10 antibody. LSM4-YFP plants were used as a positive control. Protein size is indicated.

## SUPPLEMENTARY TABLES

**Table S1.** List of enriched genes of RIP-seq of 35S:YFP-LSM4 in *lsm4-1* compared to 35S:YFP plants

**Table S2.** List of alternative splicing events altered in *lsm4-1* compared to wild-type plants

**Table S3.** List of differential expressed genes (DEG) in *lsm4-1* compared to wild-type plants

**Table S4.** List of alternative splicing events altered in *prmt5-5* compared to wild-type plants

**Table S5.** List of differentially expressed genes (DEGs) in LSM4^R^ in *lsm4-1* vs lsm4-1 plants

**Table S6.** List of differentially expressed genes (DEGs) in LSM4^RxK^ in *lsm4-1* vs *lsm4-1* plants

**Table S7.** List of alternative splicing events altered in LSM4^R^ *in lsm4-1 and lsm4-1* compared to wild-type plants

**Table S8.** List of alternative splicing events altered in LSM4^RxK^ in *lsm4-1* and *lsm4-1* compared to wild-type plants

**Table S9.** List of genes differentially splicing among LSM4^R^ and LSM4^RxK^ plants

**Table S10.** Genes affected by LSM4 methylation status that are mis-spliced upon bacterial infection by Golizs et al, 2021.

**Table S11.** List of primers used in this study

