## Supplemental Figures for "Antagonistic effects of arginine methylation of LSM4 on alternative splicing during plant stress responses"

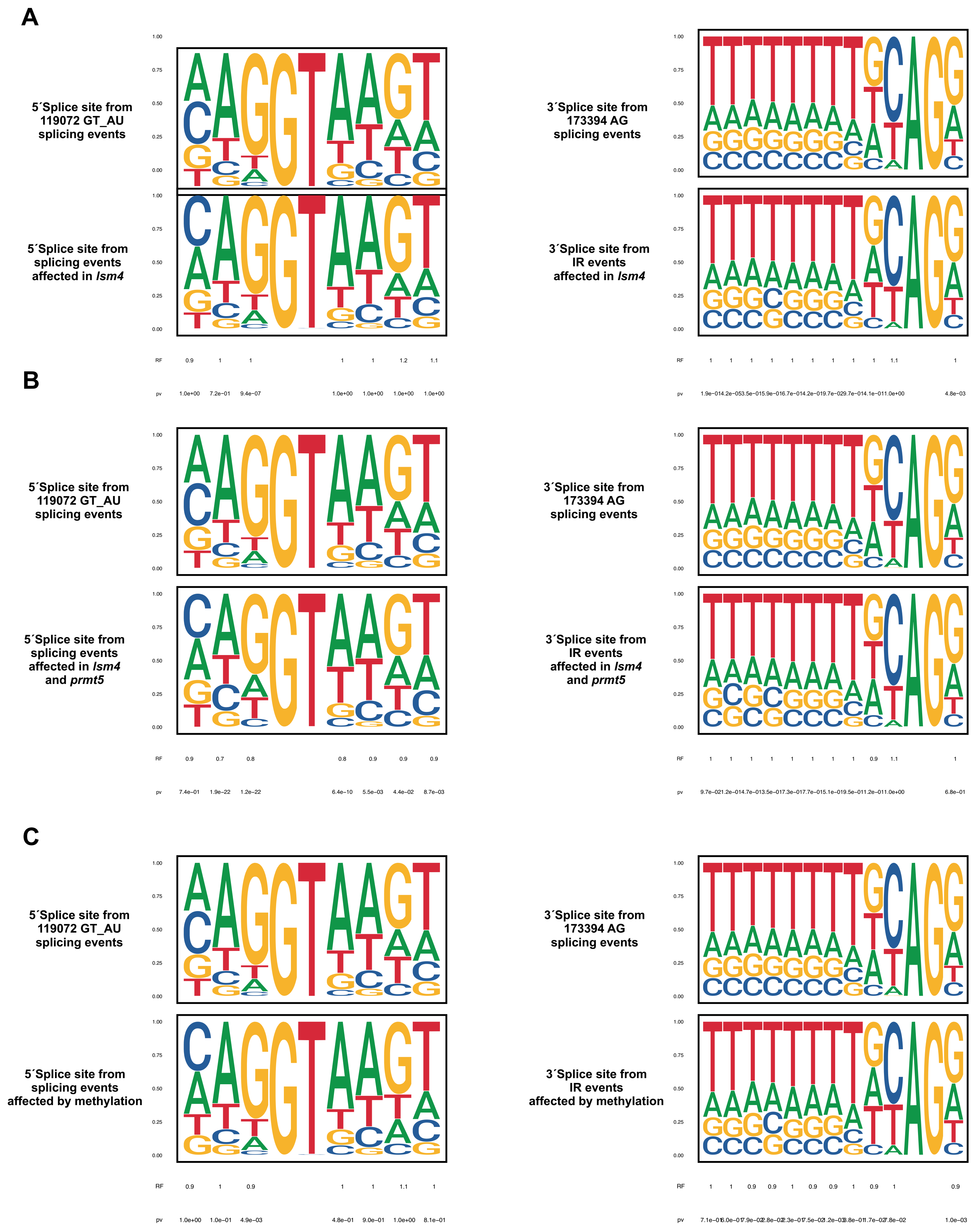

A
5´Splice site from 119072 GT_AU splicing events
5´Splice site from splicing events affected in lsm4
3´Splice site from 173394 AG splicing events
3´Splice site from IR events affected in lsm4
B
5´Splice site from 119072 GT_AU splicing events
5´Splice site from splicing events affected in lsm4 and prmt5
3´Splice site from 173394 AG splicing events
3´Splice site from IR events affected in lsm4 and prmt5
C
5´Splice site from 119072 GT_AU splicing events
5´Splice site from splicing events affected by methylation
3´Splice site from 173394 AG splicing events
3´Splice site from IR events affected by methylation
RF
pv


**Figure S1**

**A**

**LSM4 targets with DEG and DU in *lsm4-1* vs wt**

| AGI | DESCRIPTION |
|---|---|
| AT1G14250 | GDA1/CD39 nucleoside phosphatase family protein |
| AT3G44970 | Cytochrome P450 superfamily protein |
| AT4G04223 | other RNA |
| AT3G45443 | unknown protein |
| AT1G17990 | FMN-linked oxidoreductases superfamily protein |
| AT1G18020 | FMN-linked oxidoreductases superfamily protein |
| AT2G38695 | unknown protein |
| AT4G22160 | unknown protein |
| AT1G75730 | unknown protein |
| AT3G15570 | Phototropic-responsive NPH3 family protein |
| AT1G52100 | Mannose-binding lectin superfamily protein |
| AT4G24050 | NAD(P)-binding Rossmann-fold superfamily protein |
| AT3G08990 | Yippee family putative zinc-binding protein |
| AT3G13062 | Polyketide cyclase/dehydrase and lipid transport superfamily protein |
| AT3G62860 | (MAGL12) |
| AT4G11280 | 1-AMINOCYCLOPROPANE-1-CARBOXYLIC ACID (ACC) SYNTHASE 6 (ACS6) |
| AT1G33560 | ACTIVATED DISEASE RESISTANCE 1 (ADR1) |
| AT3G56970 | BHLH038 |
| AT5G12170 | CRT (CHLOROQUINE-RESISTANCE TRANSPORTER)-LIKE TRANSPORTER 3 (CLT3) |
| AT2G32190 | CYSTEINE-RICH TRANSMEMBRANE MODULE 4 (ATHCYSTM4) |
| AT4G03400 | DWARF IN LIGHT 2 (DFL2) |
| AT2G06255 | ELF4-like 3 (ELF4-L3) |
| AT1G17455 | ELF4-like 4 (ELF4-L4) |
| AT5G27720 | LSM4 |
| AT1G75220 | ERD6-LIKE 6 (ERDL6) |
| AT5G59780 | MYB DOMAIN PROTEIN 59 (MYB59) |
| AT3G61950 | MYC-TYPE TRANSCRIPTION FACTOR 67 (MYC67) |
| AT3G27660 | OLEOSIN 4 (OLEO4) |
| AT3G53350 | ROP INTERACTIVE PARTNER 3 (RIP3) |
| AT3G10450 | SERINE CARBOXYPEPTIDASE-LIKE 7 (SCPL7) |
| AT1G78290 | SNF1-RELATED PROTEIN KINASE 2-8 (SNRK2-8) |
| AT2G34070 | TRICHOME BIREFRINGENCE-LIKE 37 (TBL37) |
| AT2G28315 | UDP-XYLOSE TRANSPORTER1 (UXT1) |
| AT4G25810 | XYLOGLUCAN ENDOTRANSGLYCOSYLASE 6 (XTR6) |

**B**

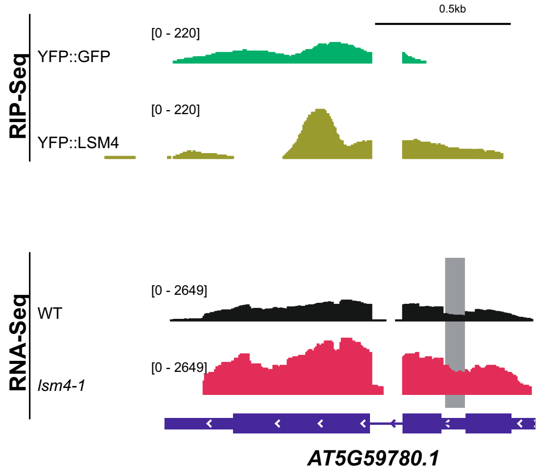


**Figure S2**

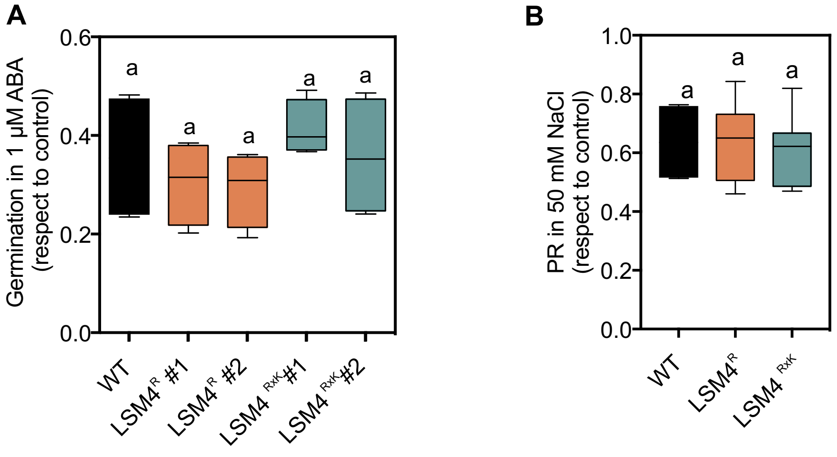
A
Germination in 1 µM ABA (respect to control)
0.6
0.4
0.2
0.0
a
a
a
a
a
WT
LSM4R #1
LSM4R #2
LSM4 RxK #1
LSM4 RxK #2
B
PR in 50 mM NaCl (respect to control)
1.0
0.8
0.6
0.4
0.2
0.0
a
a
a
WT
LSM4R
LSM4 RxK

**Figure S3**

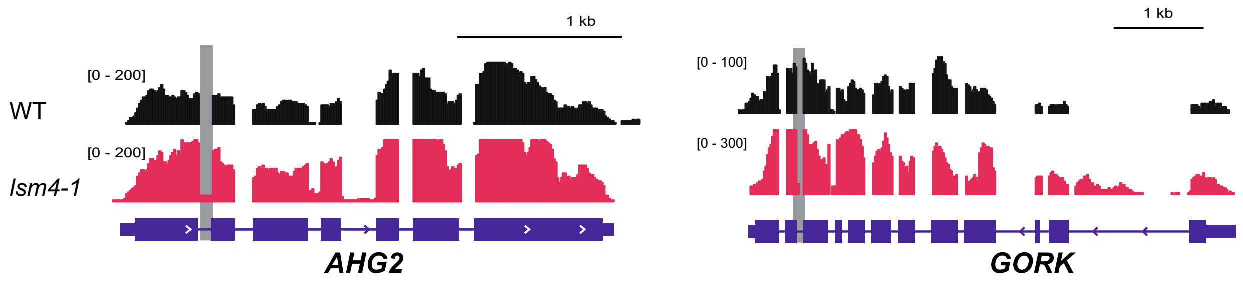


**Figure S4**

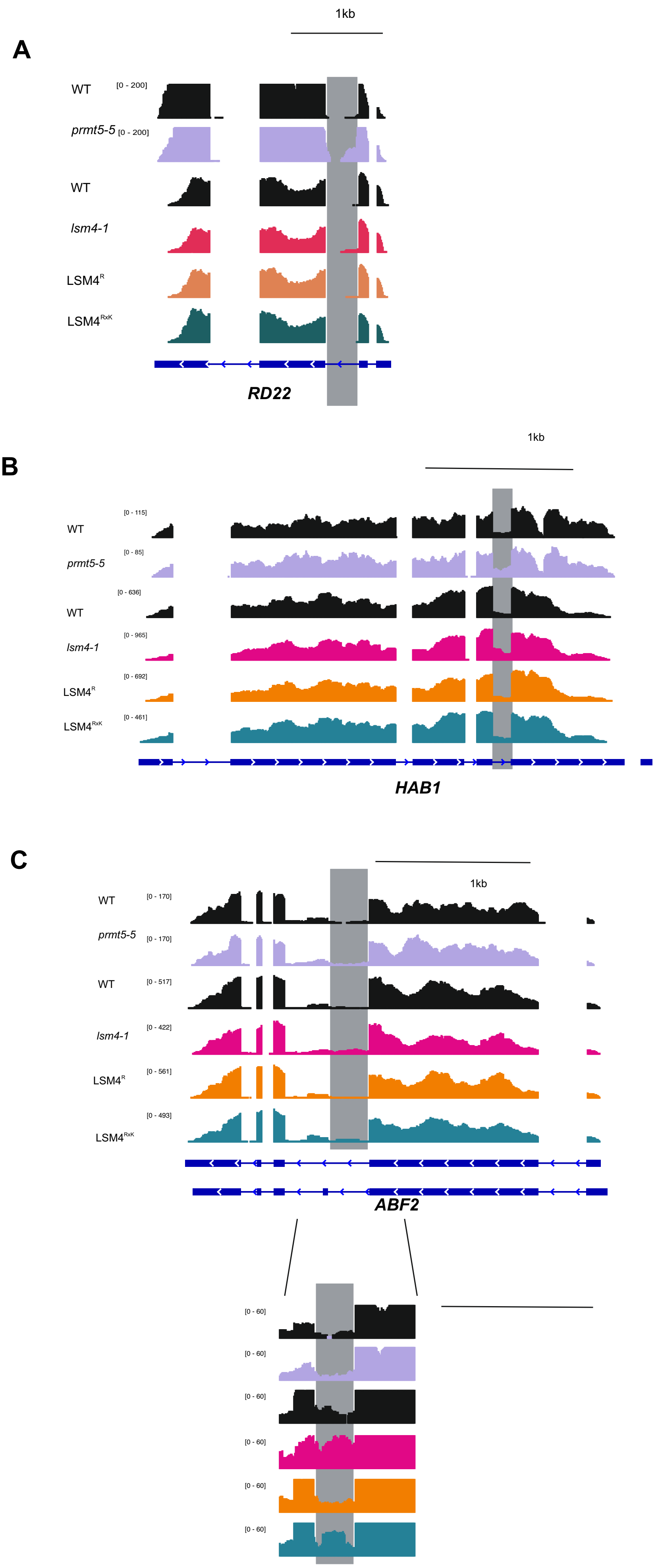

1kb
A
WT
[0 - 200]
prmt5-5
[0 - 200]
WT
lsm4-1
LSM4R
LSM4RxK
RD22
1kb
B
[0 - 115]
WT
[0 - 85]
prmt5-5
[0 - 636]
WT
[0 - 965]
lsm4-1
[0 - 692]
LSM4R
[0 - 461]
LSM4RxK
HAB1
C
1kb
WT
[0 - 170]
prmt5-5
[0 - 170]
WT
[0 - 517]
lsm4-1
[0 - 422]
LSM4R
[0 - 561]
[0 - 493]
LSM4RxK
ABF2
[0 - 60]
[0 - 60]
[0 - 60]
[0 - 60]
[0 - 60]
[0 - 60]


**Figure S5**

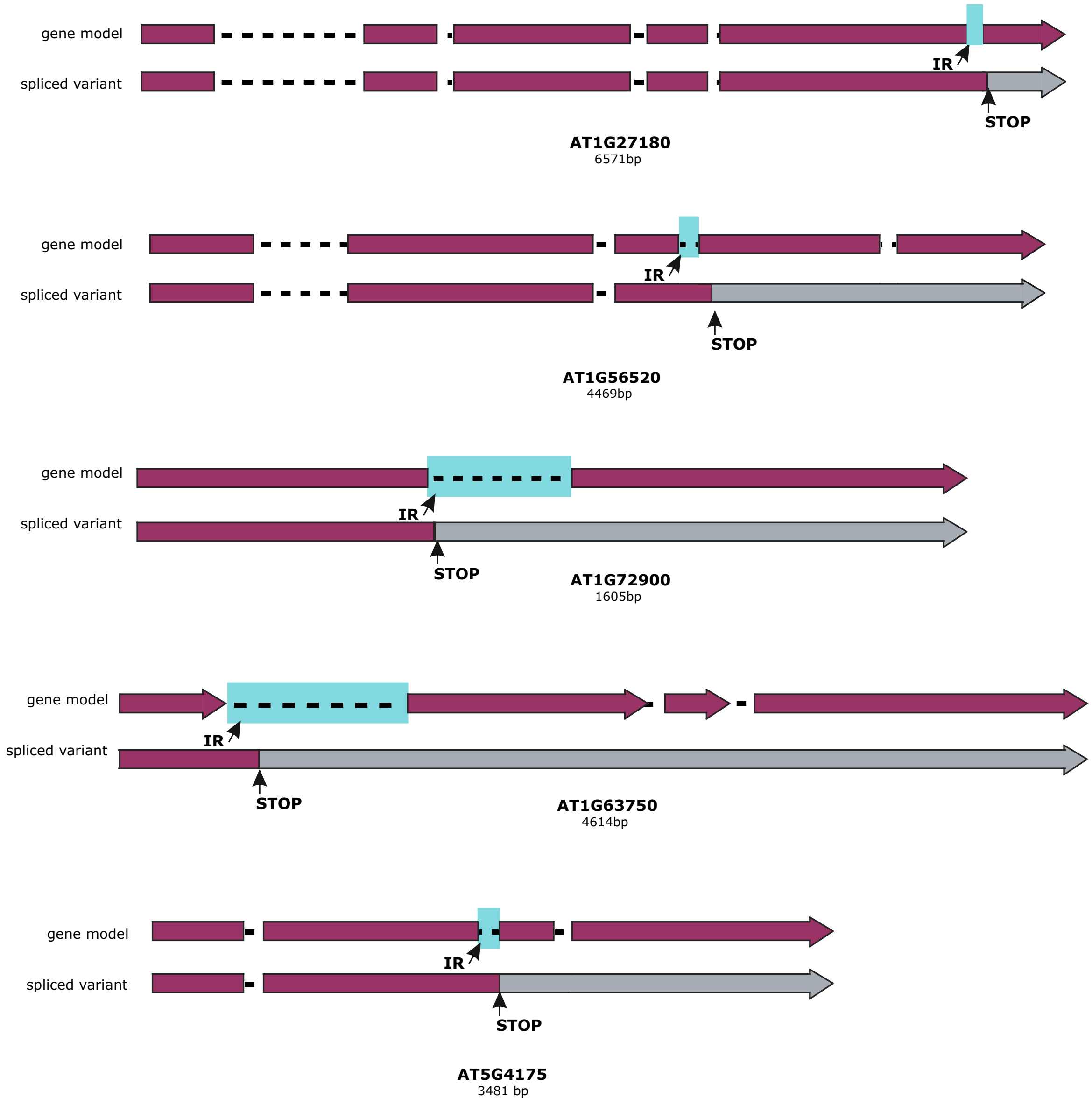
gene model
spliced variant
IR
STOP
AT1G27180
6571bp
gene model
spliced variant
IR
STOP
AT1G56520
4469bp
gene model
spliced variant
IR
STOP
AT1G72900
1605bp
gene model
spliced variant
IR
STOP
AT1G63750
4614bp
gene model
spliced variant
IR
STOP
AT5G4175
3481 bp

**Figure S6**

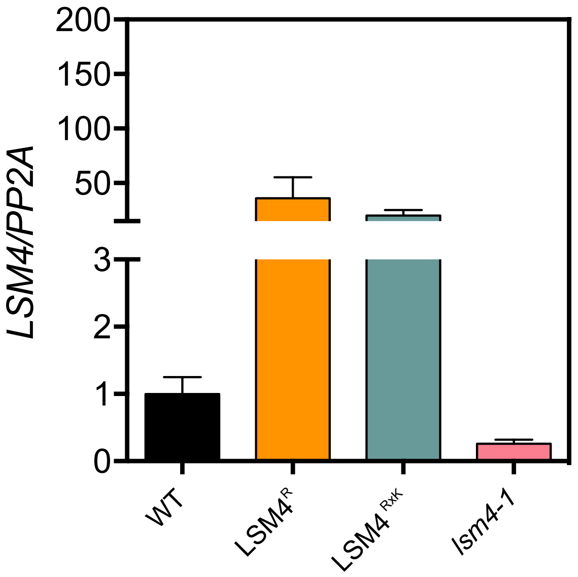


**Figure S7**

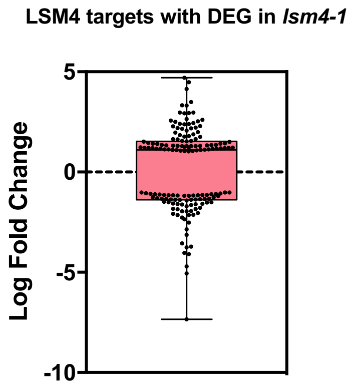


### Figure S8

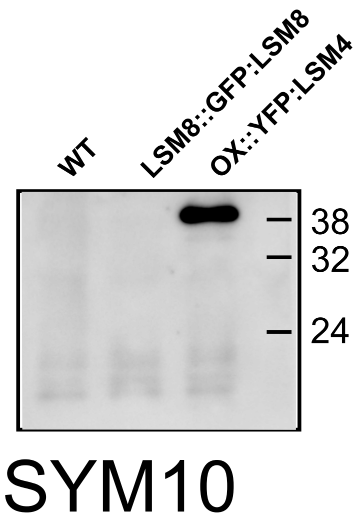
WT
LSM8::GFP:LSM8
OX::YFP:LSM4
38
32
24
SYM10

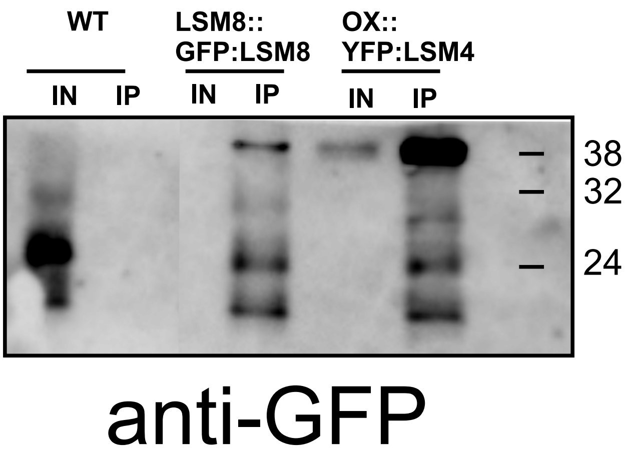
WT
LSM8::
GFP:LSM8
OX::
YFP:LSM4
IN
IP
IN
IP
IN
IP
38
32
24
anti-GFP

**Figure S9**
